# Phylogeny and chromosomal differentiation in Cestreae (Solanaceae)

**DOI:** 10.1101/2025.08.08.668976

**Authors:** Ludmila Maldonado, Jésica L. Hajduczyk Rutz, Anahi M. Yañez-Santos, Franco Chiarini, Juan D. Urdampilleta

## Abstract

Cestreae is a monophyletic tribe within the subfamily Cestroideae (Solanaceae), comprising the American genera *Cestrum*, *Sessea*, and *Vestia*. While the monophyly of *Cestrum* has been confirmed by DNA sequences, the phylogenetic relationships among its species, as well as its relationship to *Sessea* and *Vestia* remain unresolved. This study describes cytogenetic traits, expands cytological records, extends phylogenetic sampling, and analyzes chromosome evolution in Cestreae through ancestral character reconstruction. Chromosome counts, karyotype analyses, and B chromosome identification were conducted using root meristem preparations stained with Giemsa. Fluorescent in situ hybridization (FISH) with rDNA probes and DAPI was used to examine rDNA distribution. Phylogenetic relationships were inferred for 23 species using four molecular markers (*ITS, matK, ndhF,* and *trnL-F*) under Maximum Likelihood and Bayesian Inference approaches. In addition to sequences from GenBank/NCBI, 14 ITS new sequences were obtained from PCR products. Ancestral haploid chromosome numbers (*n*) were reconstructed using ChromEvol, while karyotype formulae and rDNA distribution patterns were inferred through stochastic character mapping (SCM) in *R*. All analyzed species had a chromosome number of 2*n* = 16, with two types of karyotype formula. Three *Cestrum* species possessed B chromosomes, and four distinct rDNA distribution patterns were identified. Ancestral state reconstruction suggests that 2*n* = 16, a karyotype formula of 7m + 1sm, and rDNA synteny (18-5.8-26S + 5S) on sm pair #8 and terminal 18-5.8-26S sites on one m chromosome pair, may represent ancestral characteristics in Cestreae.

## INTRODUCTION

Cestreae is a monophyletic tribe within the Cestroideae subfamily of Solanaceae, comprising three American genera, *Cestrum* L., *Sessea* Ruiz & Pav., and *Vestia* Willd. *Cestrum* comprises ca. 150 spp. distributed throughout the tropical and subtropical regions of the New World, from southern Florida and northern Mexico to Argentina (Nee, 2001). Some species, such as *Cestrum nocturnum* L., *C. elegans* (Brongn.) Schltdl., and *C. parqui* Benth., have been introduced to other regions, including Central Africa, Southeast Asia, Oceania, and several European countries. The genus *Sessea*, comprising 19 recognized species, is primarily distributed in the Andean regions of South America. Some species, such as *Sessea vestioides* (Schltdl.) Hunz. and *Sessea regnellii* Taub. are found in southern Brazil and northeastern Argentina (Benítez de Rojas & Nee, 2001; Benítez de Rojas, 2003; Keller, 2006). *Vestia* is a monotypic genus, with only *V. foetida* Hoffmanns. endemic to Chile (Hunziker, 2001). The affinity of Cestreae species is well supported by morphological, chemical, and molecular data (Faini & al., 1984; Olmstead & Palmer, 1992; Fay & al., 1998; Hunziker, 2001; Santiago-Valentin & Olmstead, 2003; Olmstead & al., 2008). *Vestia* and *Sessea* exhibit capsular fruits, whereas *Cestrum* has fleshy, berry-like fruits (Knapp, 2002). Some members of *Sessea* were included with *Cestrum* species (Carvalho & Schnoor, 1993), nevertheless, *Cestrum* is differentiated from *Sessea* by its fruit type (Francey, 1935, 1936; Benítez de Rojas & D’Arcy, 1998; Benítez de Rojas & Nee, 2001) and by molecular phylogenetic studies. Based on the characteristics of the calyx and corolla, the species of *Cestrum* were classified by Dunal (1852) in the sections *Habrothamnus* (Endl.) Schltdl. and *Eucestrum*, and Urban (1903) added the section *Pseudocestrum* to include the Hispaniolan endemic, *C. inclusum*. Some species (*C. parqui* complex, according to Nee, 2000, pers. com) are difficult to delimit due to the gradual variation of the morphological characters used in taxonomic treatments. The monophyly of *Cestrum* was confirmed with DNA sequences, but the phylogenetic relationships among its species are still unclear. Although DNA sequence data confirm the monophyly of *Cestrum*, the relationships among its species remain unresolved, despite evidence that geographical proximity can predict phylogenetic affinity within the genus (Montero-Castro & al., 2006)

The karyology of the Cestreae tribe is remarkable within Solanaceae due to several distinctive features that support its monophyly and make it an excellent model for studying karyotype evolution and the diversity of repetitive DNA sequence distribution. Key characteristics include: (1) a low chromosome number (2*n* = 16) (Tschischow, 1956; Bolkhovskikh & al., 1969; Goldblatt, 1984; Goldblatt & Johnson, 1991; Las Peñas & al., 2006); (2) large chromosome sizes (Fregonezi & al., 2006; Las Peñas & al., 2006; Fernandes & al., 2009); (3) complex heterochromatin patterns (Berg & Greilhuber, 1992, 1993; Fregonezi & al., 2006; De Paula & al., 2015); (4) the presence of B-chromosomes, or supernumerary chromosomes (Jones & Houben, 2003), in at least seven species: *Cestrum intermedium*, *C. strigilatum* (Fregonezi & al., 2004; Vanzela, 2017), *C. nocturnum* (Montechiari, 2020), *C. euanthes*, *C. parqui* (Urdampilleta & al., 2015), *C. axillare* (de Souza, 2019), and in a hybrid of *C. parqui × C. aurantiacum* (Sýkorová & al., 2003a); and finally, (5) the absence of *Arabidopsis*-type (TTTAGGG)n telomeres, which appear to be replaced by an A/T-rich minisatellite (Sýkorová & al., 2003b). On the other hand, the ribosomal DNA genes (5S and 18-5.8-26S) allow the recognition of specific or group-specific chromosomal markers, analyzing karyotypic diversity and evolutionary trends in plants (Garcia & al., 2017). Some species of Cestreae were studied using fluorescence *in situ* hybridization (FISH) to detect ribosomal DNA patterns with 5S and 18-5.8-26S probes (Fregonezi & al., 2006; Fernandes & al., 2009; De Paula & al., 2015; Urdampilleta & al., 2015). These studies detect differences in rDNA sites that suggest chromosomal rearrangements in species groups (Urdampilleta & al., 2015). This work aims (i) to describe the cytogenetic traits of 5S and 18-5.8-26S rDNA patterns in South American species of Cestreae; (ii) to expand cytological records for the tribe by providing unpublished chromosome numbers; (iii) to extend the phylogenetic sampling with these species; and (iv) to analyze chromosome numbers, rDNA patterns, and karyotypes mapped onto the molecular phylogeny obtained here, in order to reconstruct ancestral character states and observe evolutionary patterns within the tribe. In this way, cytogenetics together with molecular phylogeny will allow identifying chromosomal evolutionary trends in Solanaceae (Deanna & al., 2022).

## MATERIAL AND METHODS

### Plant material

In the present work, 19 species of the tribe *Cestreae*, 16 of the genus *Cestrum*, two of *Sessea*, and *Vestia foetida* were analyzed. Their collection data are detailed in Table 1. The new vouchers were included in the collection of the herbarium of the Botanical Museum of Córdoba (CORD).

**Table 1.**
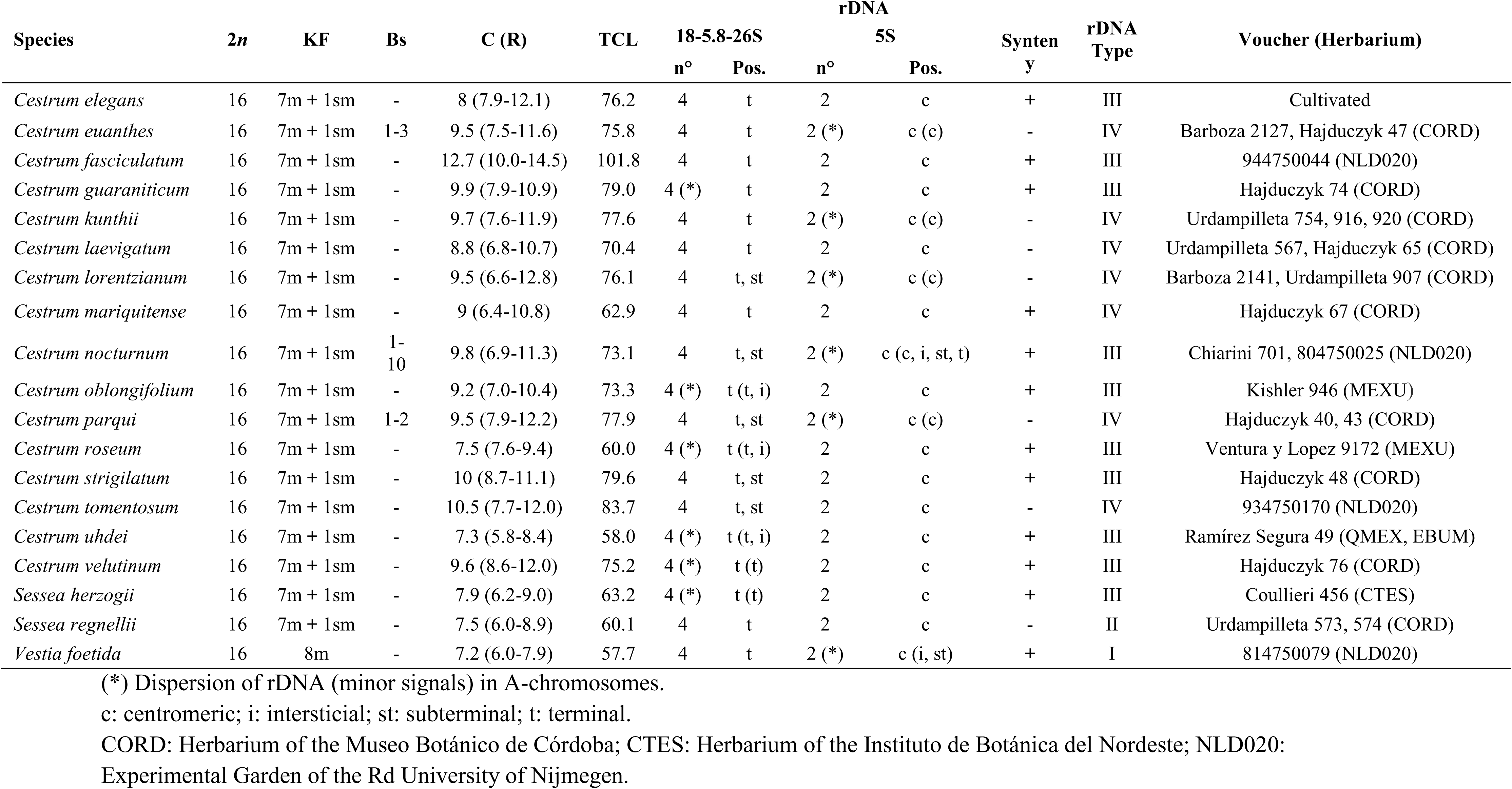
Description of the studied species of the tribe Cestreae: Chromosome number (2*n*), karyotype formula (KF), presence of B chromosomes (Bs), mean chromosome size (C) and its range (R), total chromosome length (TCL), rDNA distribution, synteny, rDNA types, and collection data of the analyzed specimens.

| Species | $2n$ | KF | Bs | C (R) | TCL | 18-5.8-26S | | rDNA<br>5S | | Synteny | rDNA<br>Type | Voucher (Herbarium) |
| --- | --- | --- | --- | --- | --- | --- | --- | --- | --- | --- | --- | --- |
|  |  |  |  |  |  | n° | Pos. | n° | Pos. |  |  |  |
| *Cestrum elegans* | 16 | 7m + 1sm | - | 8 (7.9-12.1) | 76.2 | 4 | t | 2 | c | + | III | Cultivated |
| *Cestrum euanthes* | 16 | 7m + 1sm | 1-3 | 9.5 (7.5-11.6) | 75.8 | 4 | t | 2 (*) | c (c) | - | IV | Barboza 2127, Hajduczyk 47 (CORD) |
| *Cestrum fasciculatum* | 16 | 7m + 1sm | - | 12.7 (10.0-14.5) | 101.8 | 4 | t | 2 | c | + | III | 944750044 (NLD020) |
| *Cestrum guaraniticum* | 16 | 7m + 1sm | - | 9.9 (7.9-10.9) | 79.0 | 4 (*) | t | 2 | c | + | III | Hajduczyk 74 (CORD) |
| *Cestrum kunthii* | 16 | 7m + 1sm | - | 9.7 (7.6-11.9) | 77.6 | 4 | t | 2 (*) | c (c) | - | IV | Urdampilleta 754, 916, 920 (CORD) |
| *Cestrum laevigatum* | 16 | 7m + 1sm | - | 8.8 (6.8-10.7) | 70.4 | 4 | t | 2 | c | - | IV | Urdampilleta 567, Hajduczyk 65 (CORD) |
| *Cestrum lorentzianum* | 16 | 7m + 1sm | - | 9.5 (6.6-12.8) | 76.1 | 4 | t, st | 2 (*) | c (c) | - | IV | Barboza 2141, Urdampilleta 907 (CORD) |
| *Cestrum mariquitense* | 16 | 7m + 1sm | - | 9 (6.4-10.8) | 62.9 | 4 | t | 2 | c | + | IV | Hajduczyk 67 (CORD) |
| *Cestrum nocturnum* | 16 | 7m + 1sm | 1-10 | 9.8 (6.9-11.3) | 73.1 | 4 | t, st | 2 (*) | c (c, i, st, t) | + | III | Chiarini 701, 804750025 (NLD020) |
| *Cestrum oblongifolium* | 16 | 7m + 1sm | - | 9.2 (7.0-10.4) | 73.3 | 4 (*) | t (t, i) | 2 | c | + | III | Kishler 946 (MEXU) |
| *Cestrum parqui* | 16 | 7m + 1sm | 1-2 | 9.5 (7.9-12.2) | 77.9 | 4 | t, st | 2 (*) | c (c) | - | IV | Hajduczyk 40, 43 (CORD) |
| *Cestrum roseum* | 16 | 7m + 1sm | - | 7.5 (7.6-9.4) | 60.0 | 4 (*) | t (t, i) | 2 | c | + | III | Ventura y Lopez 9172 (MEXU) |
| *Cestrum strigilatum* | 16 | 7m + 1sm | - | 10 (8.7-11.1) | 79.6 | 4 | t, st | 2 | c | + | III | Hajduczyk 48 (CORD) |
| *Cestrum tomentosum* | 16 | 7m + 1sm | - | 10.5 (7.7-12.0) | 83.7 | 4 | t, st | 2 | c | - | IV | 934750170 (NLD020) |
| *Cestrum uhdei* | 16 | 7m + 1sm | - | 7.3 (5.8-8.4) | 58.0 | 4 (*) | t (t, i) | 2 | c | + | III | Ramírez Segura 49 (QMEX, EBUM) |
| *Cestrum velutinum* | 16 | 7m + 1sm | - | 9.6 (8.6-12.0) | 75.2 | 4 (*) | t (t) | 2 | c | + | III | Hajduczyk 76 (CORD) |
| *Sessea herzogii* | 16 | 7m + 1sm | - | 7.9 (6.2-9.0) | 63.2 | 4 (*) | t (t) | 2 | c | + | III | Coullieri 456 (CTES) |
| *Sessea regnellii* | 16 | 7m + 1sm | - | 7.5 (6.0-8.9) | 60.1 | 4 | t | 2 | c | - | II | Urdampilleta 573, 574 (CORD) |
| *Vestia foetida* | 16 | 8m | - | 7.2 (6.0-7.9) | 57.7 | 4 | t | 2 (*) | c (i, st) | + | I | 814750079 (NLD020) |

(\*) Dispersion of rDNA (minor signals) in A-chromosomes.
c: centromeric; i: interstitial; st: subterminal; t: terminal.
CORD: Herbarium of the Museo Botánico de Córdoba; CTES: Herbarium of the Instituto de Botánica del Nordeste; NLD020:
Experimental Garden of the Rd University of Nijmegen.

### Cytogenetic techniques

The chromosome preparations were obtained from root meristems pretreated with 2mM 8-hydroxyquinoline for 4-5h at 14°C, fixed in ethanol:acetic acid (3:1, v:v) for 24h at room temperature and stored at -20°C until use. The tissues were digested with Pectinex enzyme solution (Novozimes) and squashed in 45% acetic acid. Preparations were frozen in liquid nitrogen to remove the coverslip. The preparations were immersed in a 2% solution of Giemsa for conventional staining and mounted in Entellan (Merck).

The fluorescent in situ hybridization (FISH) was developed using the methodology described by Schwarzacher & Heslop-Harrison (2000). The probe p*Ta*71 has been used to map the rDNA loci in 18S-5.8S-26S (Gerlach & Bedbrook, 1979), labeled with biotin-14-dUTP by nick translation (Bionick, Invitrogen) and subsequently detected with avidin-FITC (Sigma). For the analysis of 5S rDNA loci, a probe was obtained by PCR amplification using 5L1 and 5L2 primers (Shibata & Hizume, 2002), using genomic DNA of *C. parqui* L’Her. as a template. The 5S rDNA fragments were labeled with Digoxigenin-11-dUTP (DIG Nick translation mix, Roche) and detected with Anti-DIG-Rhodamine (Roche). Post-hybridization washing was performed with 2x SSC, 0.1xSSC, 2× SSC, and 4x SSC/0.2% Tween, 10 min, 42°C each, and the slides were mounted with antifade Vectashield (Vector Laboratories), containing 1.5 ug/ml of DAPI (4′,6-diamidino-2-phenylindole).

The metaphases were photographed using an Olympus BX61 microscope coupled with a monochromatic camera and Cytovision software (Leica Biosystems) for documentation. The chromosomes were organized and measured using Photoshop CS4 software (Adobe Systems Inc.) and MicroMeasure v3.3 (Reeves, 2001). The chromosomes were classified (*m*, *sm*, *st*, or *t*) and designated according to Levan & al. (1964).

### Molecular markers

For this study, four molecular markers were used: the nuclear ribosomal internal transcribed spacer (ITS), and three plastid markers, the maturase K gene (*mat*K), the NADH dehydrogenase subunit F gene (*ndh*F), and the intron of the *trn*L gene added to the intergenic spacer between the *trn*L and *trn*F genes (*trn*LF) (Supplementary Table 1). For ITS, 14 new sequences were obtained from sequencing of the PCR product. The remaining sequences used come from the database available in Genbank/NCBI (Supplementary Table 1).

To obtain ITS sequences, DNA was isolated from leaf tissue using the CTAB II protocol (Weising & al., 2005). ITS regions were amplified via PCR using the ITS4 and ITS5 primers (White & al., 1990). PCR amplifications were performed in a MasterCycler (Eppendorf) with a total reaction volume of 25 μL, containing 1.5 mM MgCl2, 0.2 μM of each primer, 0.2 mM dNTPs, and 0.025 U/μL of GoTaq DNA polymerase (Promega). Amplification products were verified by electrophoresis on a 1% agarose gel, stained with SYBR Safe (Invitrogen, Eugene, OR, USA), and visualized using a UV transilluminator. The PCR products were sequenced by Macrogen using the Sanger method with the same primers.

### Alignment, Phylogenetic Analysis and Ancestral States Reconstruction (ASR)

The sequences were edited and aligned with MEGA 11 (Tamura & al., 2021) using the iterative MUSCLE method and concatenated with SequencesMatrix (Vaidya & al., 2011). A phylogenetic analysis was performed using four species, *Browallia americana* L., *B. speciosa* Hook., *Streptosolen jamesonii* (Benth.) Miers and *Salpiglossis sinuata* Ruiz & Pav. as outgroups (Olmstead & al., 2008) (Supplementary Table 1). For the maximum likelihood analysis (ML) the best substitution model was chosen by jModelTest 2.1.4 (Darriba & al., 2012) using Corrected Akaike’s Information Criterion (AICc). The analyses were performed in raxmlGUI 2.0 (Edler & al., 2021) using the GTR + Γ model with bootstrapping (1.000 replicates). Although both jModelTest and raxmlGUI 2.0 suggested the TVM+Γ4 model, the GTR + Γ model was chosen based on availability in the software and the difference in AICc was not significant (Supplementary Table 2). Bayesian analysis (BI) was conducted with MrBayes 3.2 (Ronquist & al., 2012) using the same substitution model, with 2 runs, 4 Markov chains Monte Carlo (MCMC) for 10.000.000 generations, retaining one tree each every 1.000th generation, and a burnin of 25%. The convergence was checked with Tracer 1.7 (Rambaut & al., 2018). To obtain the MCC maximum credibility tree, the TreeAnnotator tool belonging to the BEAST software package (Suchard & al., 2018) was used, which required a concatenation of the runs performed in MrBayes. This concatenation was carried out through the R v4.3.1 (R Core Team, 2023), with package *ape* (Paradis & al., 2004). To compare the phylogenetic trees obtained with ML and BI, we used R v4.3.1 (R Core Team, 2023), with the packages *ape* (Paradis & al., 2004), *phytools* (Revell, 2012), *TreeDist* (Smith, 2020) and *phangorn* (Schliep, 2019) to graph and calculate the Robinson-Foulds distance (Robinson & Foulds, 1981).

For ASR we ultrametricized the MCC tree using the *chronos* function from the R packages *ape* (Paradis et al., 2004) and *phytools* (Revell, 2012). The ancestral haploid chromosome numbers (n) were reconstructed in ChromEvol (Glick & Mayrose, 2014). All models obtained were tested using AIC to select the best fitting model (Akaike, 1974). We set the maximal chromosome number as –10 and the minimal chromosomal number as 1 (according to ChromEvol settings), and 10.000 simulations. To perform an optimized ChromEvol run, a model comparison was performed, from which the CONST_RATE model was selected for having the lowest AIC (Supplementary Table 3). The chromosome numbers of *B. americana*, *B. speciosa, S. jamesonii,* and *S. sinuata* were obtained from CCDB (Rice & Mayrose, 2023).

For karyotype formula and rDNA distribution patterns (18S-5.8S-26S and 5S), we performed a maximum likelihood reconstruction of ancestral states as a function of stochastic character mapping (SCM) (Huelsenbeck & al., 2003) in R v4.3.1 (R Core Team, 2023), with packages *ape* (Paradis & al., 2004), *geiger* (Harmon & al., 2008) and *phytools* (Revell, 2012). The ER (equal transition rates), SYM (forward and reverse transitions share the same parameter), and ARD (all transition rates are different) models were fitted for the karyotype formula and rDNA distribution patterns. For this, the *fitDiscrete* function of package *geiger* was used, and the best model was selected according to AIC criterion. The best-fitting model for character transformation was the ER model (Supplementary Table 4). The ancestral states reconstruction was carried out using maximum likelihood, with the Markov k-state (Mk1) model according to the better-adjusted transition model.

## RESULTS

### Chromosome number and karyotype structure

The chromosome number of *C. guaraniticum* Chodat & Hassl., *C. oblongifolium* Schltdl., *C. roseum* Kunth, *C. uhdei* Dammer ex Francey, *C. velutinum* Hiern and *S. herzogii* Dammer is reported for the first time. The number 2*n* = 16 was found in all species studied, and B-chromosomes (supernumerary chromosomes) were observed in *C. elegans* (Brongn.) Schltdl., *C. nocturnum* L., *C. parqui* Benth (Table 1, Fig. 1). Typically, the karyotypes were highly symmetrical, having seven m (#1-7) plus one submetacentric pairs (#8), except for *V. foetida* Hoffmanns which has eight metacentric (*m*) pairs. The A-chromosome complement ranged from approx. of 6 to 13 μm, total chromosome length (TCL) of 58 to 79 μm, and an average chromosome length of 7.2 to 9.8 μm. The B-chromosome were 5-6 times smaller, ranging from 1.7 to 2.8 μm (Table 1).

**Figure 1.**
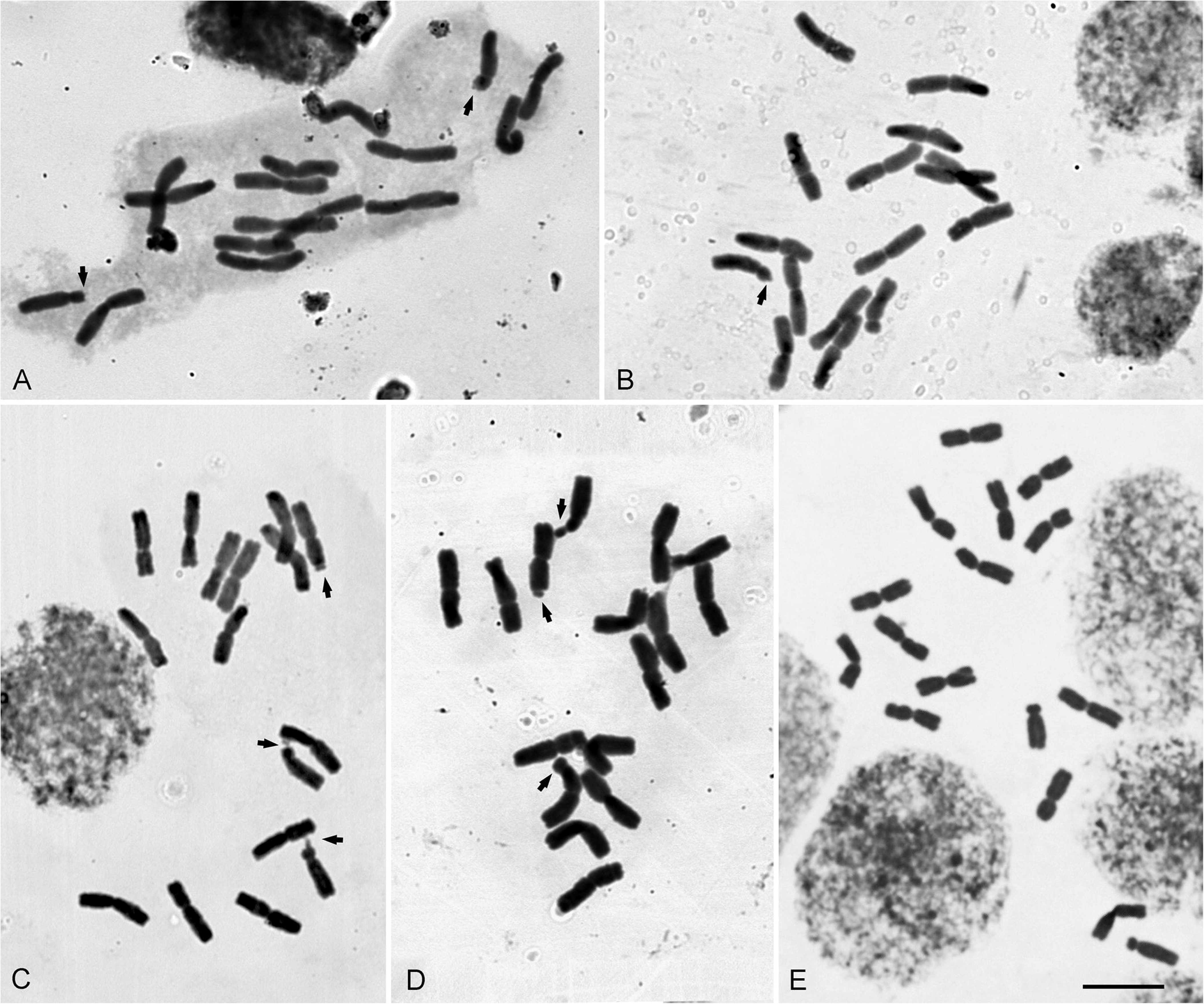
Mitotic metaphase chromosomes stained by the conventional technique (Giemsa) in Cestreae species with new count. Arrows indicate secondary constrictions. **A.** *C. guaraniticum*; **B.** *C. oblongifolium*; **C.** *C. roseum*; **D.** *C. uhdei*; **E.** *S. herzogii*. Bar = 10 µm.

### FISH and Chromosomal distribution of rDNA genes

Following FISH analysis, five species presented two different types of 18-5.8-26S rDNA signals according to size with six species exhibiting both major and minor 5S rDNA signals (Table 1, Fig. 2). The number of major 18-5.8-26S and 5S rDNA loci was conserved, being typically four sites for 18-5.8-26S and two for the 5S rDNA. Due to possible rDNA dispersion, the loci number of minor 18-5.8-26S and 5S rDNA is variable in some species. In *C. oblongifolium*, *C. roseum, C. uhdei,* and *C. velutinum* up to 6 minor 18-5.8-26S sites were observed. Several minor hybridization signals of 5S rDNA were detected in variable numbers in *C. norturnum* and *V. foetida,* and one minor signal of 5S rDNA in *C. euanthes, C. kunthii*, *C. lorentzianum,* and *C. parqui*. Intercalar, subterminal, and terminal DAPI bands (AT rich heterochromatin) were detected with variation between most species (Fig. 2).

**Figure 2.**
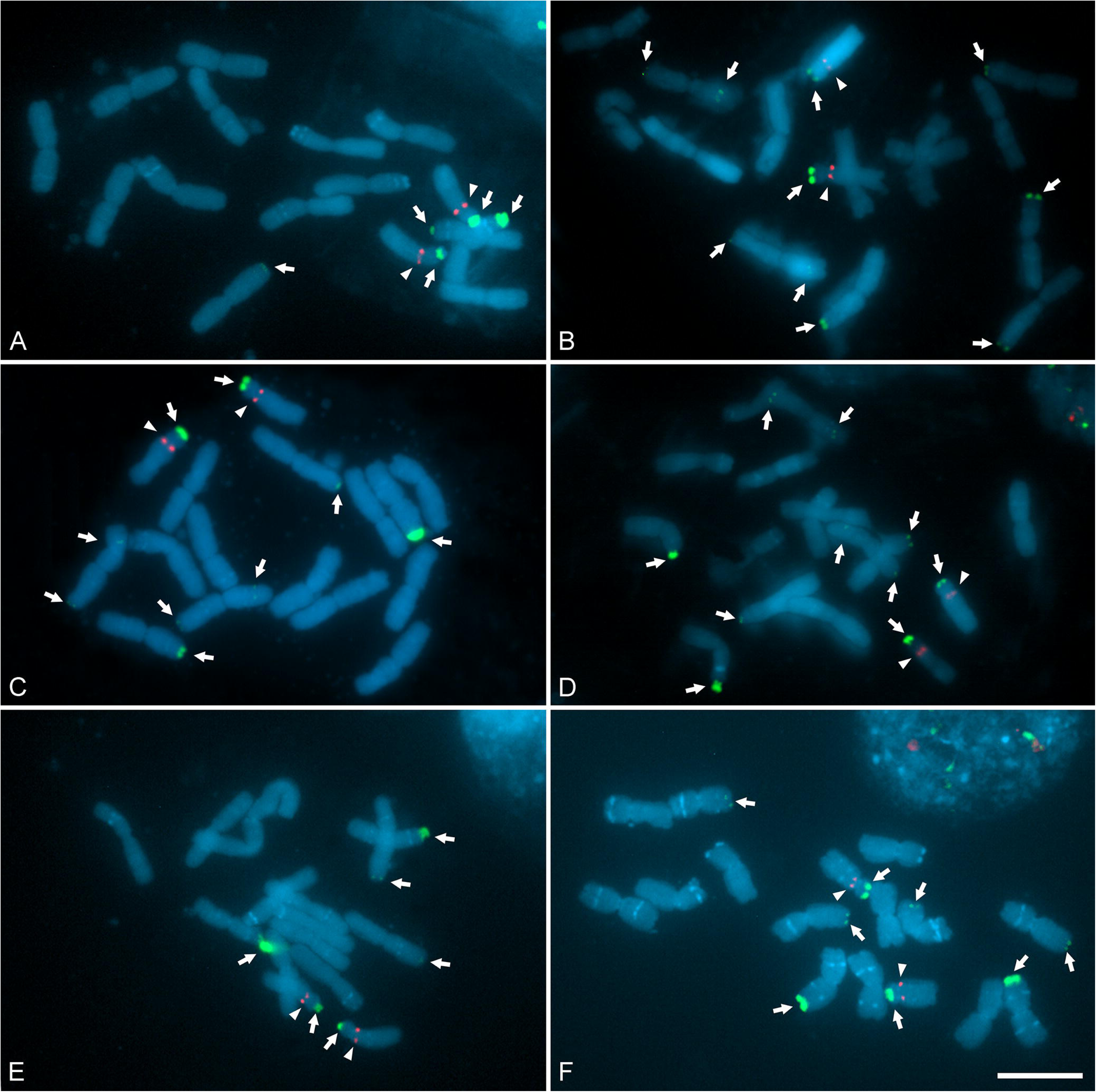
Distribution of rDNA loci (18-5.8-26S, arrow, green, and 5S, arrowhead, red) in Cestreae. **A.** *C. guaraniticum*. **B.** *C. oblongifolium*. **C.** *C. roseum*. **D.** *C. uhdei*. **E.** *C. velutinum* **F.** *S. herzogii*. Bar = 10 µm.

The position of major 18-5.8-26S and 5S rDNA loci was variable among some species and allowed us to recognize four general patterns of chromosomal distribution (I, II, III, and IV types) (Table 1, Fig. 2, 3). Type I was only observed in *V. foetida*, which was characterized by showing two pairs of 18-5.8-26S rDNA hybridization signals in terminal regions of *m* chromosomes, and one of them has a pair of 5S rDNA signals in the pericentromeric region. Additionally, minor 5S rDNA hybridization signals were observed in various chromosomes, together with four terminal and two intercalar pairs of DAPI+ bands. Type II was only observed in *S. regnellii*, with two pairs of 18-5.8-26S rDNA sites in terminal regions of *m* and *sm* (#8) chromosomes and a 5S rDNA locus in the region pericentromeric of other *m* chromosome pairs. Also, *S. regnellii* showed no DAPI bands in terminal or intercalary regions. The Type III pattern observed is characterized by colocalization of 18-5.8-26S and 5S rDNA in the *sm* chromosome pair (#8) (rDNA synteny) and one pair of 18-5.8-26S rDNA sites in the terminal region of a *m* chromosome. This pattern is common in Cestreae and was detected in *C. elegans*, *C. fasciculatum*, *C. guaraniticum*, *C. mariquitense, C. laevigatum*, *C. nocturnum*, *C. oblongifolium*, *C. roseum*, *C. strigilatum*, *C. uhdei*, *C. velutinum*, *S. herzogii* (Table 2, Fig. 2, 3). This group is characterized by a high amount of terminal, subterminal, and intercalar DAPI+ bands, frequently heteromorphic (as *C. strigilatum* and *C. nocturnum*). Type IV was observed in the *C. parqui* complex (*C. euanthes*, *C. lorentzianum*, *C. parqui), C. kunthii,* and *C. tomentosum*, with two pairs of intense 18-5.8-26S rDNA signals in two *m* pairs, and one pair of 5S rDNA sites in pericentromeric regions of a *sm* chromosome pair (#8). Subterminal to terminal and minor intercalary DAPI+ bands were detected in some pairs of chromosomes, in *C. lorentzianum*, *C. parqui,* and *C. tomentosum* terminal DAPI+ bands and distal to the 18-5.8-26S rDNA sites were observed (Table 1, Fig. 2, 3). Because we do not have information on the distribution of rDNA genes of the outgroup species, we have assigned them the "Other" category.

**Figure 3.**
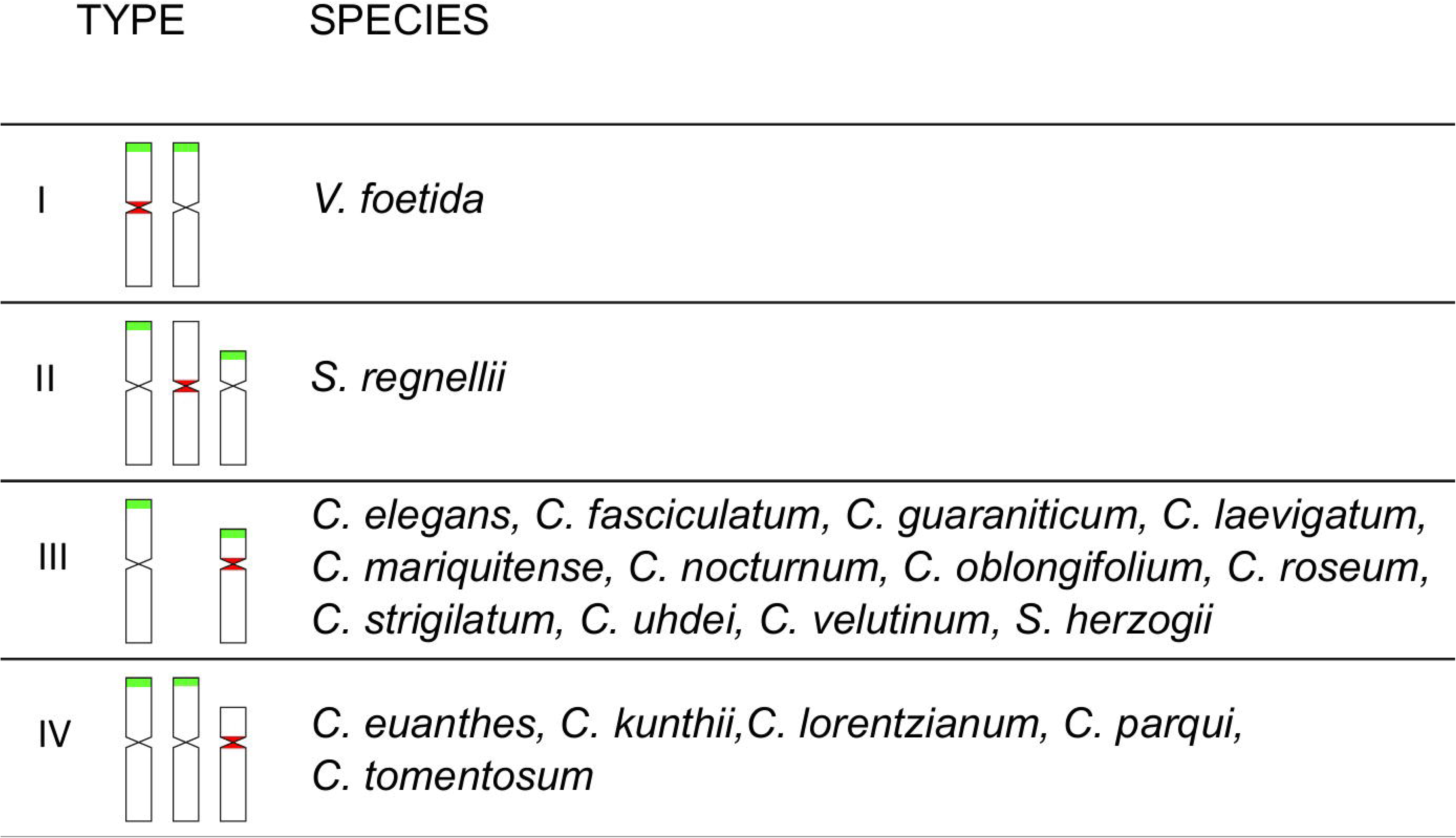
Scheme of the four types of major rDNA loci organization (18-5.8-26S, green, and 5S, red) observed in Cestreae.

**Table 2.** Description of the matrix of aligned sequences, considering of the four molecular markers (ITS, *mat*K, *ndh*F, *trn*LF) their terminal numbers, the alignment size, their position in the matrix, C-G richness which is expressed as a percentage, as well as the number of conserved, variable and informative sites.

|  | **ITS** | ***matK*** | ***ndhF*** | ***trnLF*** | **Concatenated** |
| --- | --- | --- | --- | --- | --- |
| **Number of terminals** | 24 | 13 | 14 | 14 | 24 |
| **Alignment size** | 719 | 1888 | 2108 | 1040 | 5755 |
| **Position in alignment** | 1-719 | 720-2607 | 2608-4715 | 4716-5755 | 1-5755 |
| **Richness CG (%)** | 59.2 | 32.6 | 33.3 | 34.0 | 36.4 |
| **Conserved sites** | 512 | 1670 | 1842 | 841 | 4865 |
| **Variable sites** | 204 | 132 | 188 | 94 | 618 |
| **Parsim-Info sites** | 114 | 38 | 95 | 25 | 272 |

### Phylogenetic relationships with *ITS*, matK, ndhF and trnLF

The phylogeny of Cestreae is inferred with 24 tips (23 species), adding new information on 14 ITS sequences from 13 species. The complete ITS region of all samples (24 sequences from 23 species), the *mat*K gene (13 sequences from 13 species), the NADH dehydrogenase subunit F gene (14 sequences from 14 species), and the *trn*LF region (14 sequences from 14 species) were analyzed and are summarized in Table 2. The sequence matrix had 5755 bp that includes ITS (1-719), *mat*K (720-2607), *ndh*F (2608-4715), and *trn*LF (4716-5755). On average, the richness of CG bases in these regions is almost 40%. The sequences showed relative variability. Of all bases, 4865 bases were conserved characters, while 618 and 272 were respectively variable and informative characters. In particular, the ITS region generated varies from 683 to 709 bp, having the alignment 719 bp, with 114 (15,86%) parsimoniously informative sites. These are reduced to 26 when considering only the parsimony informative sites found among species of the genus *Cestrum*. The *mat*K gene varies from 765 to 1794 bp in the alignment of 1888 bp, with 38 (2,01%) parsimoniously informative sites. The NADH dehydrogenase subunit F gene varies from 1046 to 2108 bp in the alignment of 2108 bp, with 95 (4,51%) parsimoniously informative sites. Finally, the *trn*LF region varies from 286 to 946 bp in the alignment of 1040 bp, with 25 (2,40%) parsimoniously informative sites. Table 2 describes the general characteristics of the matrices included in the phylogenetic analysis, and the alignment and resulting trees are archived on TreeBase (accessions ID 31997). The log likelihood of the best tree with ML analysis is - 12296.59 (Fig. 4A) and showed a similar topology with MCC tree (BI) (Fig. 4B). The Robinson-Foulds distance (Robinson & Foulds, 1981) calculated for the trees obtained by ML and BI was 8. This measure, together with the comparison of the trees (Fig. 4), shows us that there are no major inconsistencies. As a complementary approach, maximum likelihood (ML) and Bayesian inference (BI) analyses were conducted separately for the nuclear (ITS) and plastid (*mat*K, *ndh*F, and *trn*LF) markers, to verify potential incongruence between the results derived from the two genomes. The main differences may be associated with the limited plastid data, as these reconstructions were based on 15 species, compared to the 23 species (including two samples of *C. nocturnum*) used in the ITS analyses (Supplementary Figures S1–S6).

**Figure 4.**
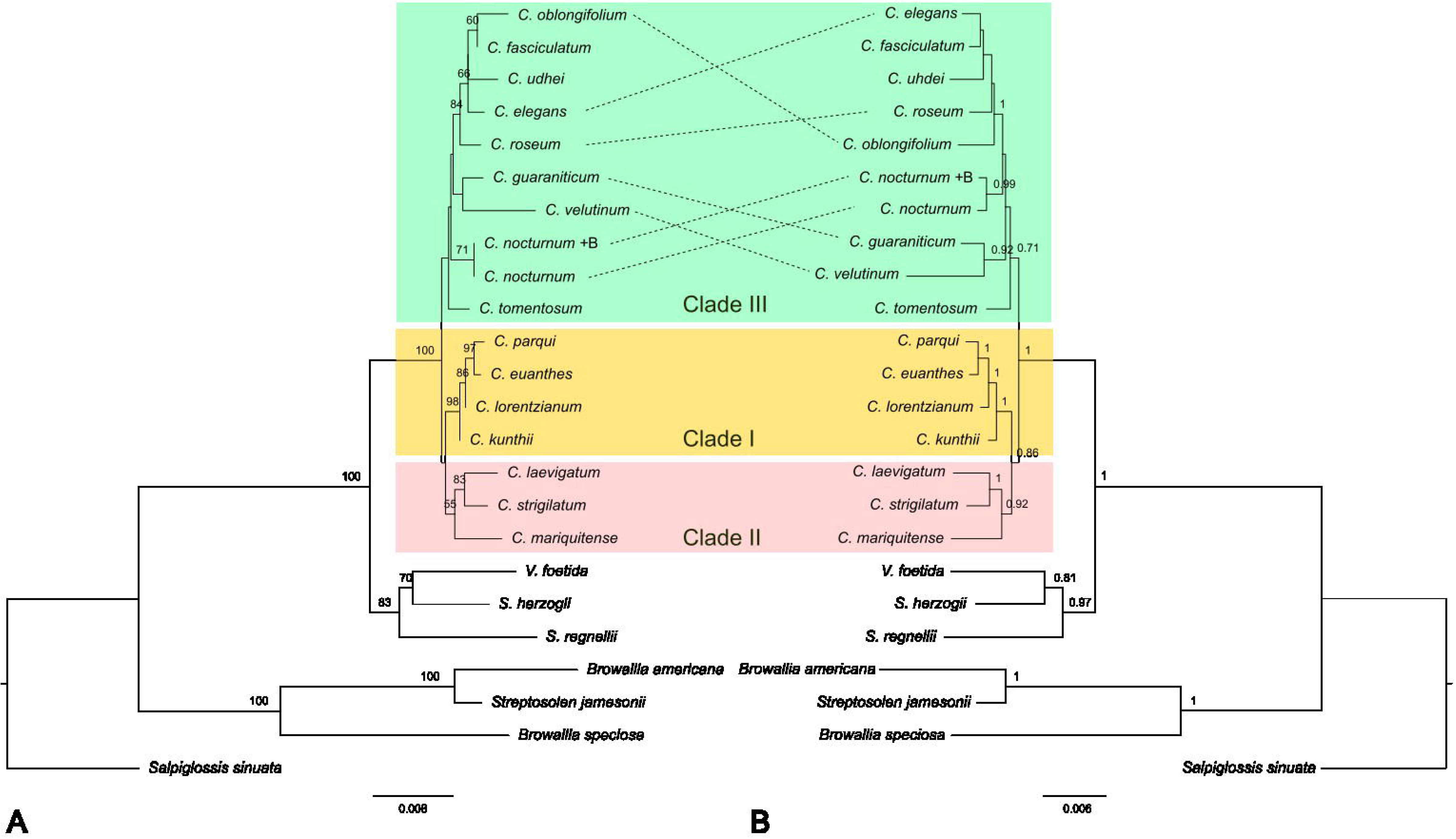
Comparison of phylogenetic relationships among the analyzed Cestreae species, based on the molecular markers ITS, *mat*K, *ndh*F and *trn*LF, using *Browalia americana, B. speciosa, Streptosolen jamesonii* and *Salpiglossis sinuata* as outgroups. *Cestrum nocturnum* +B indicates the presence of B chromosomes. In yellow, Clade I is highlighted, in pink the Clade II and in green is the Clade III. **A.** ML analysis (-12296.59) with bootstraps greater than 50. **B.** BI analysis with posterior probabilities greater than 0.5.

Regarding the phylogenetic relationships between species, we observed that *Cestrum* forms a well-supported monophyletic clade, while *Sessea* does not appear to be monophyletic, as *Vestia* is nested within it; however, this may become clearer by including additional *Sessea* species and incorporating more molecular markers or phenotypic characters (Fig. 4). Despite some polytomies due to little variation in DNA sequences, *Cestrum* species analyzed form three clades, Clade I: *C. euanthes*, *C. lorentzianum*, *C. kunthii* and *C. parqui;* Clade II: *C. laevigatum, C. mariquitense* and *C. strigilatum;* Clade III: *C. elegans*, *C. fasciculatum*, *C. oblongifolium*, *C. roseum, C. uhdei, C. nocturnum, C. velutinum, C. guaraniticum* and *C. tomentosum* (Fig. 4). ITS sequences were obtained in *C. nocturnum*, both from plants with and without B chromosomes, the sequences being identical in base pairs.

### Ancestral States Reconstruction of chromosome characters

The Cestreae species have base chromosome numbers of x = 8, and all species analyzed were diploid with 2*n* = 16. The ML analysis (ChromEvol 2.0), which was based on the MCC tree (BI), enabled the ancestral chromosome base number for the common ancestor of Cestreae tribe to be inferred as x = 8 (pp = 0.95) (Fig. 5).

**Figure 5.**
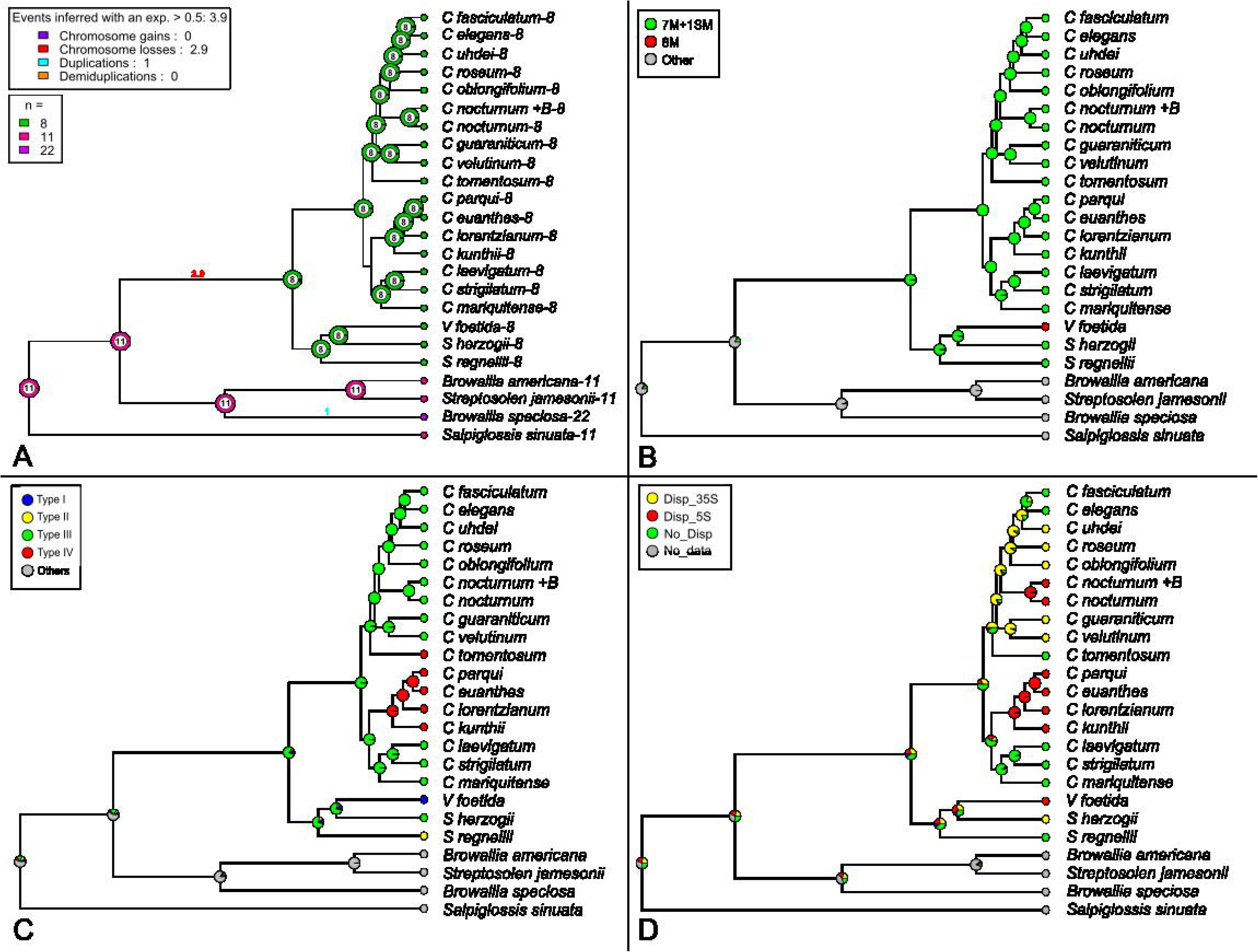
Ancestral state reconstruction of chromosome features in Cestreae and related taxa on the MCC tree. **A.** Analysis of base chromosome numbers with ChromEvol. **B.** Mapping using the maximum likelihood reconstruction methods of karyotype formula. **C.** 18-5.8-26S and 5S rDNA distribution patterns. **D.** rDNA dispersal events.

The karyotype formula and localization of rDNA loci obtained were mapped on the BI phylogenetic tree using the maximum likelihood reconstruction methods. The ER model was the one that best fitted to the chromosomal traits studied here (Supplementary Table 4). The analyses resulted in the ASR for the karyotype formula and 18-5.8-26S and 5S rDNA distribution suggests that the ancestral state of the tribe Cestreae is most likely 7m + 1sm and had a Type III chromosome rDNA distribution pattern, including two terminal 18-5.8-26S rDNA loci and one pericentromeric 5S rDNA, without synteny between both rDNA types (Fig. 5). On the other hand, the ASR concerning rDNA implies that some dispersal events arose independently in different Cestreae groups. The minor rDNA 18-5.8-26S sites occur in *C. oblongifolium*, *C. roseum*, *C. uhdei*, *C. velutinum* and *S. herzogii.* The minor rDNA 5S sites were observed in *C. euanthes*, *C. lorentzianum*, *C. kunthii*, *C. parqui, C. nocturnum,* and *V. foetida* (Fig. 5).

## DISCUSSION

### Phylogeny of tribe Cestreae

Although the morphological diversity within the tribe Cestreae is remarkable, the molecular divergence in the *mat*K, *ndh*F, *trnL*F, and the nuclear ITS region is relatively limited. Specifically, the ITS sequences have shown a higher percentage of informative characters (15%) compared to previous estimates (4% and 2%). These findings align with earlier studies suggesting a recent and rapid diversification of the tribe (Montero-Castro & al., 2006). Our analysis of these sequences confirms that Cestreae is a monophyletic group, with *Sessea* forming a clade distinct from *Cestrum*. The relationship between *Sessea* and *Vestia* would require validation. Although both the molecular data obtained in this study and the shared fruit type (capsule) suggest the inclusion of *V. foetida* within the genus *Sessea*, we consider it necessary to conduct a more comprehensive analysis that includes a greater number of species, additional molecular markers and/or morphological variables. This is important because there are relevant differences that should be considered, such as seed morphology: in *Sessea* species, seeds are winged, whereas in *Vestia* they lack wings and are more similar to those of *Cestrum* (Hunziker, 2001).

Although some species of the genus *Sessea* were transferred to *Cestrum* sect. *Sessea* (Carvalho & Schnoor, 1993), this perspective was not fully accepted (Benítez de Rojas & Nee, 2001). The molecular markers used in this study, along with rDNA distribution patterns, clearly distinguish *Sessea/Vestia* from *Cestrum* (Fig. 6). This is consistent with Knapp (2002), who suggested that the soft, indehiscent fruits (berries) of *Cestrum* likely evolved independently from the capsular fruits of *Vestia* and *Sessea*, which are more common in Solanaceae.

**Figure 6.**
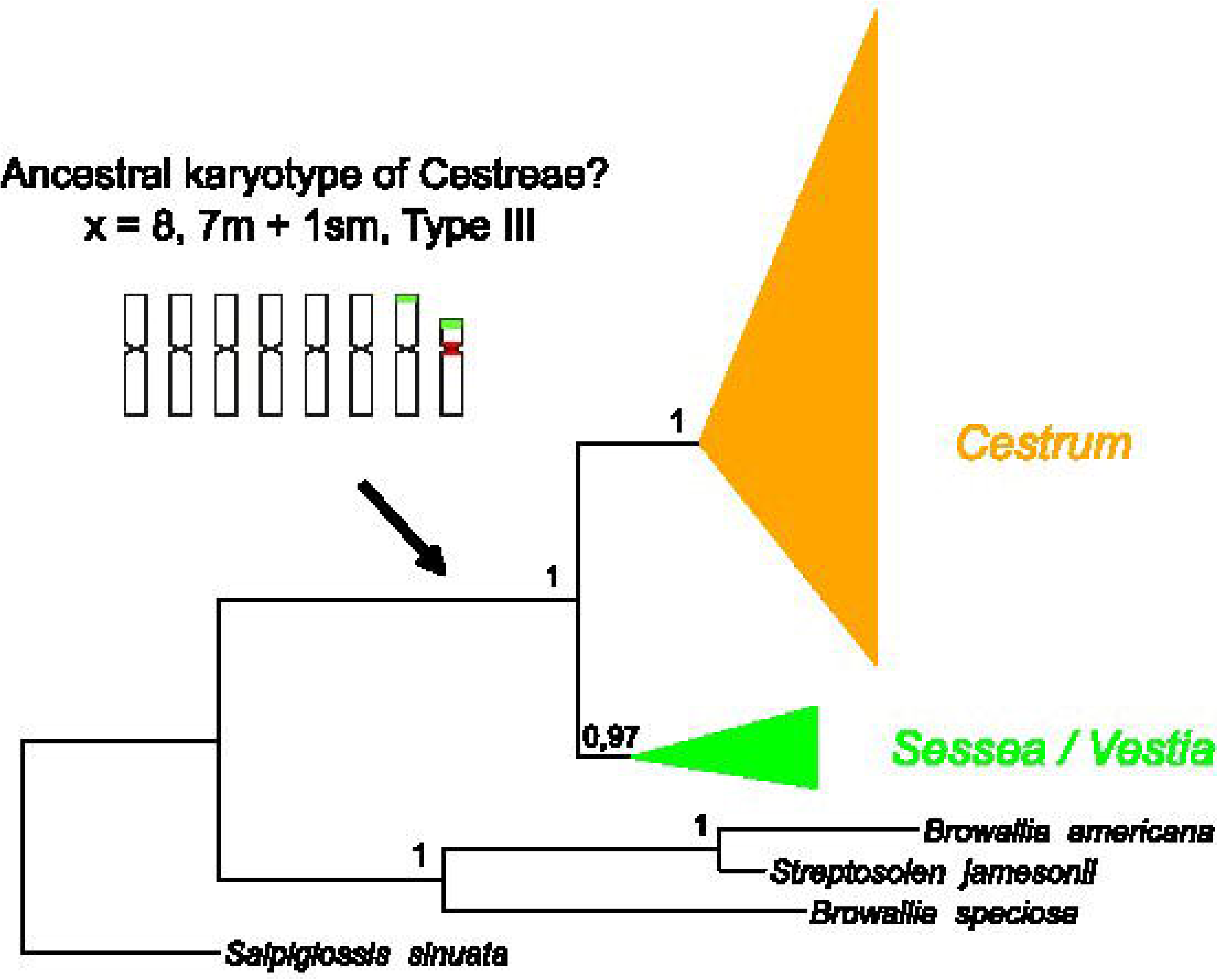
Result of the reconstruction of the ancestor of the tribe Cestreae.

The present study aligns in certain aspects with the phylogeny presented by Montero-Castro & al. (2006) in considering the genus *Cestrum* as monophyletic. Both studies reveal geographically structured clades and do not support the monophyly of the sections proposed by Dunal (1852) and Urban (1903). This is likely because these classifications are mainly based on floral structures associated with pollination syndromes, which may have evolved convergently without reflecting true phylogenetic relationships. In this context, the differences between the two studies lie in the species included. Specifically, this study incorporated South American species that form clades I and II. Regarding clade III, *C. uhdei*, *C. velutinum*, and *C. guaraniticum* were newly included, while *C. elegans*, *C. fasciculatum*, *C. oblongifolium*, and *C. roseum* were grouped by Montero-Castro within the "Central-Mex" clade. On the other hand, *C. nocturnum* was placed in the "Guatemala" clade, and *C. tomentosum* in a sister clade to the "Hispaniola" clade. We consider that including more species from Mexico, Guatemala, Costa Rica, Panama, and the Caribbean could improve the resolution of clade III. However, phylogenetic relationships still require further resolution due to differences in sampling, the use of different molecular markers, and the low molecular divergence between species, consistent with recent diversification.

### Evolution of Chromosome number in Cestreae

The results confirm the chromosome numbers previously reported for other species of Cestreae (Tschischow, 1956; Sharma & Sharma, 1959; Bolkhovskikh & al., 1969; Moscone, 1992; Fregonezi & al., 2006; Las Peñas & al., 2006; Fernandes & al., 2009; Urdampilleta & al., 2015). In this work are presented the first report for *C. guaraniticum*, *C. oblongifolium*, *C. roseum*, *C. uhdei*, *C. velutinum*, *S. herzogii*. The species studied in this work were diploid with 2*n* = 16, and these results support that the basic number (x = 8) is highly conserved within the tribe Cestreae. However, only 10-15% of the species of the tribe have been analyzed for their chromosome number. The haploid number n = 8 is also shared by other genera, such as *Bouchetia* DC. ex Dunal, *Hunzikeria* D’Arcy and *Nierembergia* Ruiz & Pav. However, this character could be considered homoplastic within Solanaceae (Deanna & al., 2022). Despite this, the differentiation between the subfamilies Cestroideae and Solanoideae may be associated with variation in the basic chromosome number. Chromosome counts based on x = 11 are common in genera of the subfamily Cestroideae, as reported in *Browallia*, *Streptosolen* and *Salpiglossis* (Moscone, 1992; Badr & al., 1997). Although the chromosome number of *Protoschwenkia* Soler. remains unknown, the data from the subfamily Cestroideae suggest a reduction from x = 11 to x = 8 in species of the tribe Cestreae. This hypothesis is strongly supported by different phylogenetic hypotheses and the reconstruction of the ancestry of the chromosome number. The ancestral chromosome number of the subfamily Cestroideae is likely x = 11, and the reduction to x = 8 likely occurred before the diversification of the tribe Cestreae (ca. 10 My) (Särkinen & al., 2013).

### B chromosomes in Cestrum

The number of chromosomes within the cell can vary due to the presence of small accessory chromosomes, the B chromosomes. The B chromosomes had been previously reported in some *Cestrum* species (Sýkorová & al., 2003b; Fregonezi & al., 2004; Urdampilleta & al., 2015). These chromosomes vary between and within species through processes of meiotic and mitotic instability, respectively (Montechiari & al., 2020). In carrier species, the Bs are 3-5 folds smaller than A-chromosomes, as usually reported in plants (Jones & Houben, 2003). The B chromosomes reported in *Cestrum* species exhibit a similar morphology; however, variations in heterochromatin patterns and the distribution of repetitive DNA have been reported (Fregonezi & al., 2004; Urdampilleta & al., 2015; Montechiari & al., 2020). The B chromosomes are not present in all *Cestrum* species and are found in different clades, therefore, the common vs. independent origin has not yet been clarified.

### Karyotype variation in Cestreae

The species of tribe Cestreae are characterized by large chromosomes and symmetrical highly karyotypes (Fregonezi & al., 2006; Las Peñas & al., 2006; Fernandes & al., 2009). The majority of species analyzed have seven metacentric pairs (*m*) and one submetacentric pair (*sm*). The constancy of chromosome shape and size between species in some Solanaceae taxa was used as evidence of karyotypic orthoselection in the family, which preserves rather similar complements because they seem to be more stable (Moscone & al., 2003). However, the conservation of chromosome shape and number in Cestreae could simply be due to the recent diversification of the species (Särkinen & al., 2013; Huang & al., 2023).

Due to the presence of a karyotype with 8m in *V. foetida*, which reflects a more symmetric karyotype within Cestreae, this characteristic was considered ancestral (Las Peñas & al., 2006). However, our preliminary results based on ASR suggest that *V. foetida* is closely related to species of the genus *Sessea*, and that a karyotype with 7m + 1sm may have existed in the common ancestor of Cestreae, reflecting a trend towards increasing karyotypic symmetry in some species. Nonetheless, the phylogenetic relationship between *V. foetida* and species of the genus *Sessea* still requires clarification, which could be achieved through broader sampling.

### Diversity patterns of rDNA distribution and evolutionary trends

The rDNA distribution of Cestreae allows us to characterize some groups of species according to relative localization of 5S and 18-5.8-26S rDNA loci. Our results confirm previous works on *Cestreae* (Fregonezi & al., 2006; Fernandes & al., 2009; De Paula & al., 2015; Urdampilleta & al., 2015) and report new data on the distribution of 18-5.8-26S and 5S rDNA sites in *C. guaraniticum, C. oblongifolium, C. roseum, C. uhdei, C. velutinum, S. herzogii*. All species studied of Cestreae have conserved the terminal and pericentromeric positions for 18-5.8-26S and 5S rDNA, respectively, principally for major sites. Despite the constancy of the karyotype features and numbers of major rDNA loci, the Cestreae genera and *Cestrum* groups studied for the moment can be differentiated by the relative distribution of these 18-5.8-26S and 5S rDNA sites (Fig. 3), and grouped in four different types (I, II, III, and IV). These results contribute to the knowledge of karyotype diversity in the group and the value of rDNA distribution as a cytotaxonomic character in Cestreae could be complemented with species of the tribe not yet studied, especially the little-known species of *Sessea*.

The ancestral character reconstruction suggests that the distribution of type III rDNA is present in different clades of Cestreae, including *S. herzogii* and several species of *Cestrum*. This characteristic could be present in the common ancestor of Cestreae, and the loss of colocalization of rDNA sites (18-5.8-26S and 5S rDNA) could be characteristics derived for this group. If this hypothesis were correct, the Type I, II, and IV distribution could be caused by different rearrangements as chromosome translocations and equilocal dispersion of repeat DNA (Roa & Guerra, 2015). The Type I distribution in *Vestia* involves the transformation of pair #8 into a metacentric chromosome, maintaining the synteny of the 18-5.8-26S and 5S rDNA loci on one of the metacentric chromosomes. The Type II distribution in *S. regnelli* maintains the submetacentric #8 pair, but it is the only species known so far of Cestreae that possesses a pericentromeric 5S rDNA site on a metacentric chromosome. The type IV distribution present in *C. euanthes*, *C. kunthii*, *C. parqui*, *C. lorentzianum*, could be caused by chromosomal rearrangements that lead to the loss of synteny of the rDNA loci in chromosome pair #8, maintaining the rDNA regions 5S in pericentromeric position.

The presence and dispersion of rDNA minor sites could be recognized as a characteristic derived in this group. The presence of minor rDNA sites has also been observed in other species of Solanaceae, such as those in the genus *Capsicum* (Romero-da Cruz & al., 2017), *Solanum* (Moyetta & al. 2017; Mesquita & al., 2024), *Nolana* and *Sclerophylax* (Lujea & Chiarini, 2017). Different mechanisms have been postulated to explain the mobility of rDNA sites (Raskina & al., 2008; Lan & Albert, 2011). Similar to what occurs in other plants, the dispersion of 5S and 18-5.8-26S rDNA in Cestreae may have occurred independently across different lineages, utilizing distinct mobility mechanisms for repeat DNA while preserving an equilocal distribution pattern.

## CONCLUSION

In summary, our results confirm the monophyly of the tribe Cestreae and identify two distinct clades: one comprising *Cestrum* species and the other containing *Sessea* with the monotypic *Vestia* nested inside. Additionally, within the genus *Cestrum*, two new clades are recognized that include South American species. The molecular divergence in the ITS, *mat*K, *ndh*F, and *trn*LF regions is relatively low, suggesting rapid diversification, which poses challenges for accurately resolving phylogenetic relationships. Ancestral state reconstruction indicates that 2*n* = 16, a karyotype formula of 7m + 1sm, and Type III rDNA distribution could represent ancestral characteristics in Cestreae (Fig. 6). Further studies incorporating additional species, particularly from the genus *Sessea*, are needed to resolve the position of *Vestia* with respect to *Sessea* as currently circumscribed. However, it would also be valuable to include species from more geographically diverse regions. Another promising approach could involve incorporating a greater number of molecular markers.

## ACKNOWLEDGEMENTS

The authors are grateful to the Argentinean agencies: CONICET, ANPCyT-FONCyT, MINCyT-Córdoba, and SECyT-Universidad Nacional de Córdoba for financial support. They also thank the reviewers and the associate editor for their valuable suggestions to improve the manuscript.

## AUTHOR CONTRIBUTIONS

LM performed the bioinformatics analyses, wrote and edited the manuscript and edited the figures. JLHR performed the scientific experiments and data collection. AMYS performed the scientific experiments and reviewed the manuscript. FC performed the scientific experiments, contributed to the identification of the plant material, data collection and reviewed the manuscript. JDU designed, performed and supervised the scientific experiments, data collection, wrote and reviewed the manuscript and edited the figures. All authors contributed to the article and approved the submitted version.

## APPENDICES LEGENDS

**Supplementary Table 1.** GenBank sequences.

**Supplementary Table 2.** AIC and delta AIC values obtained from model comparison (jModelTest) to be used in ML analysis.

**Supplementary Table 3.** AIC values obtained from the comparison of models used for the optimization of the ChromEvol.

**Supplementary Table 4.** AIC values obtained from the comparison of models used to Reconstruct the Ancestral States of the karyotypic formula and the rDNA distribution.

**Supplementary Figures (S1-S6).** Phylogenetic analyses using maximum likelihood (ML) and Bayesian inference (BI) conducted with nuclear markers (S1 and S2), plastid markers (S3 and S4) and the combined dataset (S5 and S6).

## Notes

### Competing Interest Statement

The authors have declared no competing interest.

