## Supplementary Figures for "Phylogeny and chromosomal differentiation in Cestreae (Solanaceae)"

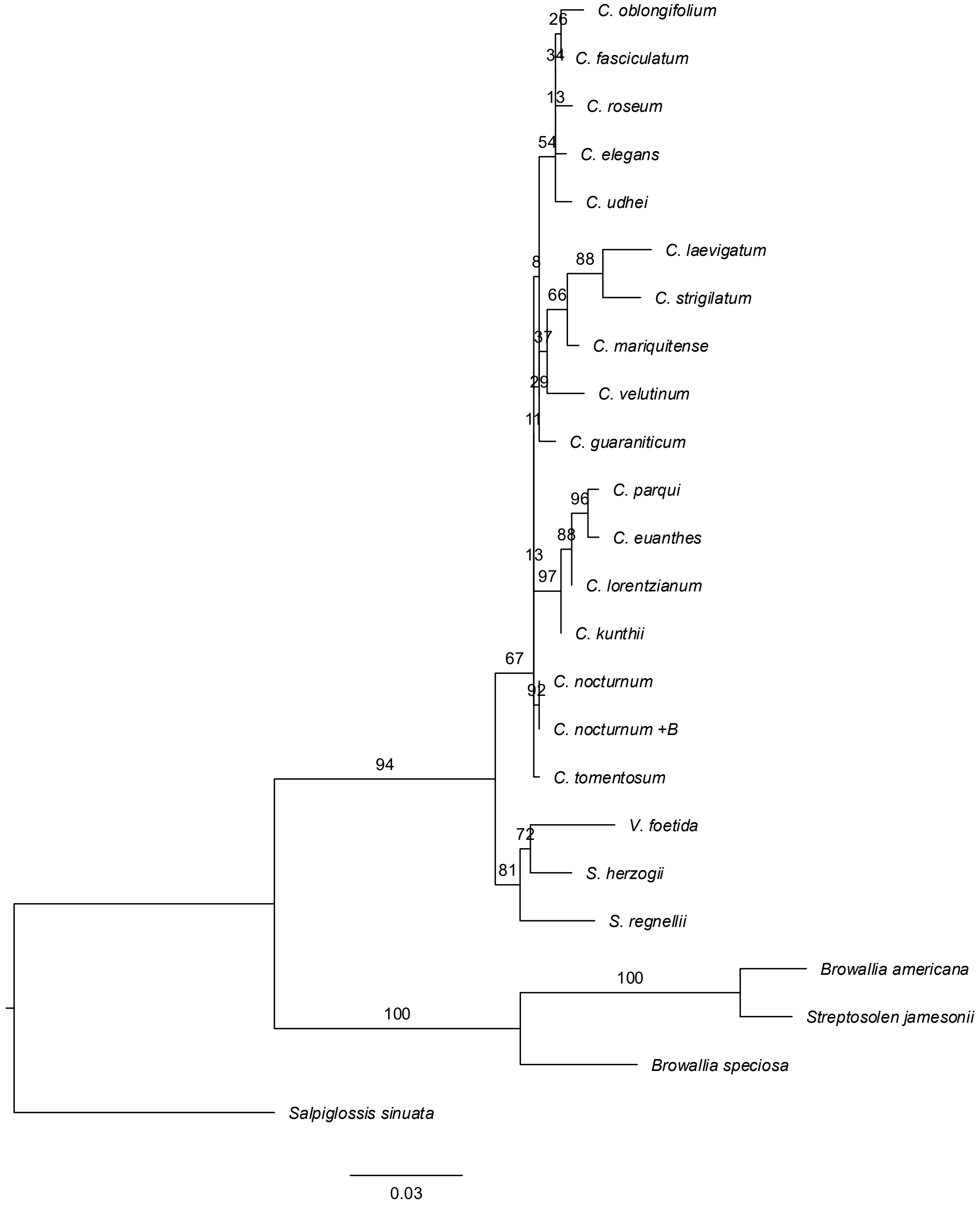


Supplementary Figure S1: ML analysis with ITS.

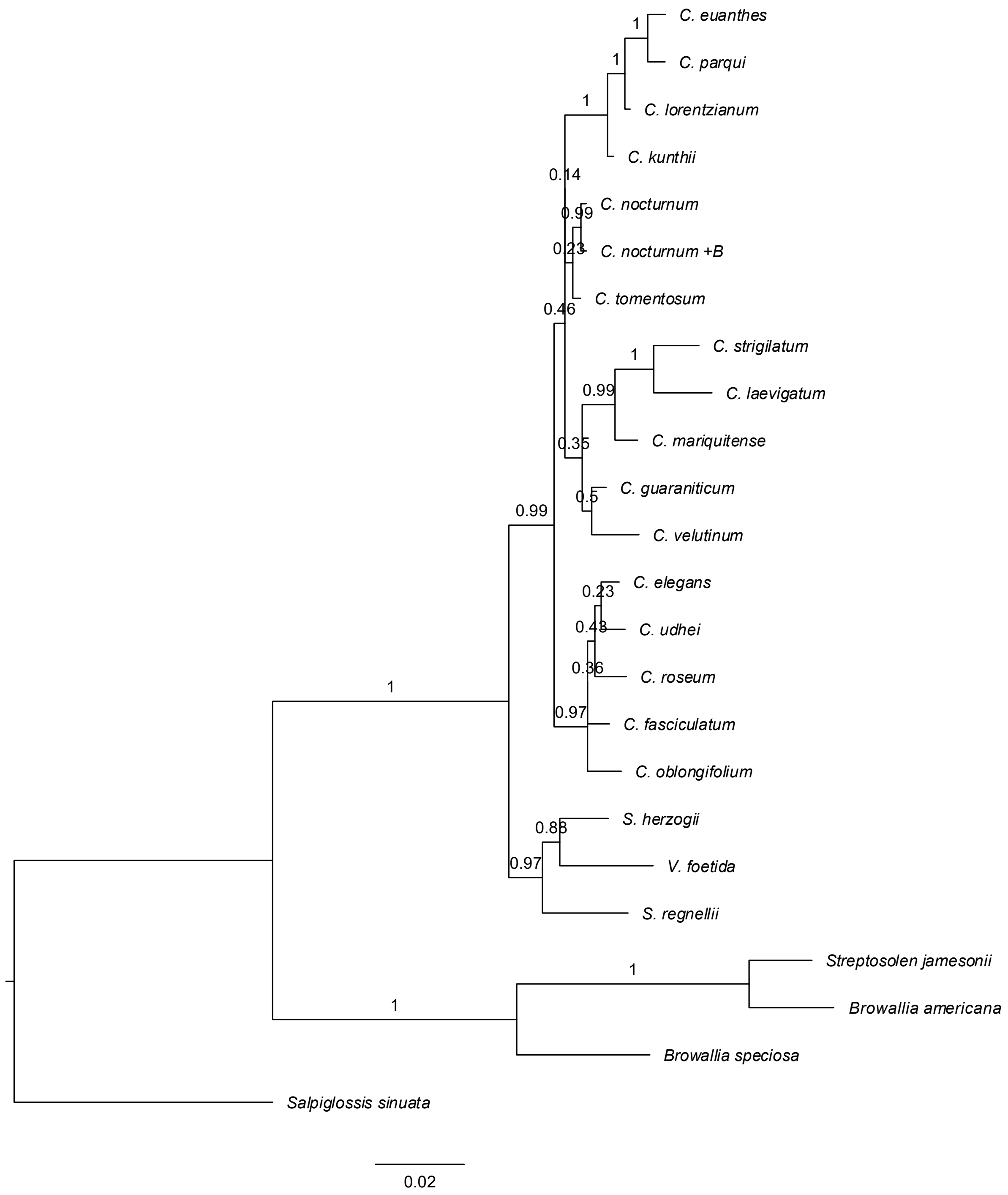


Supplementary Figure S2: BI analysis with ITS.

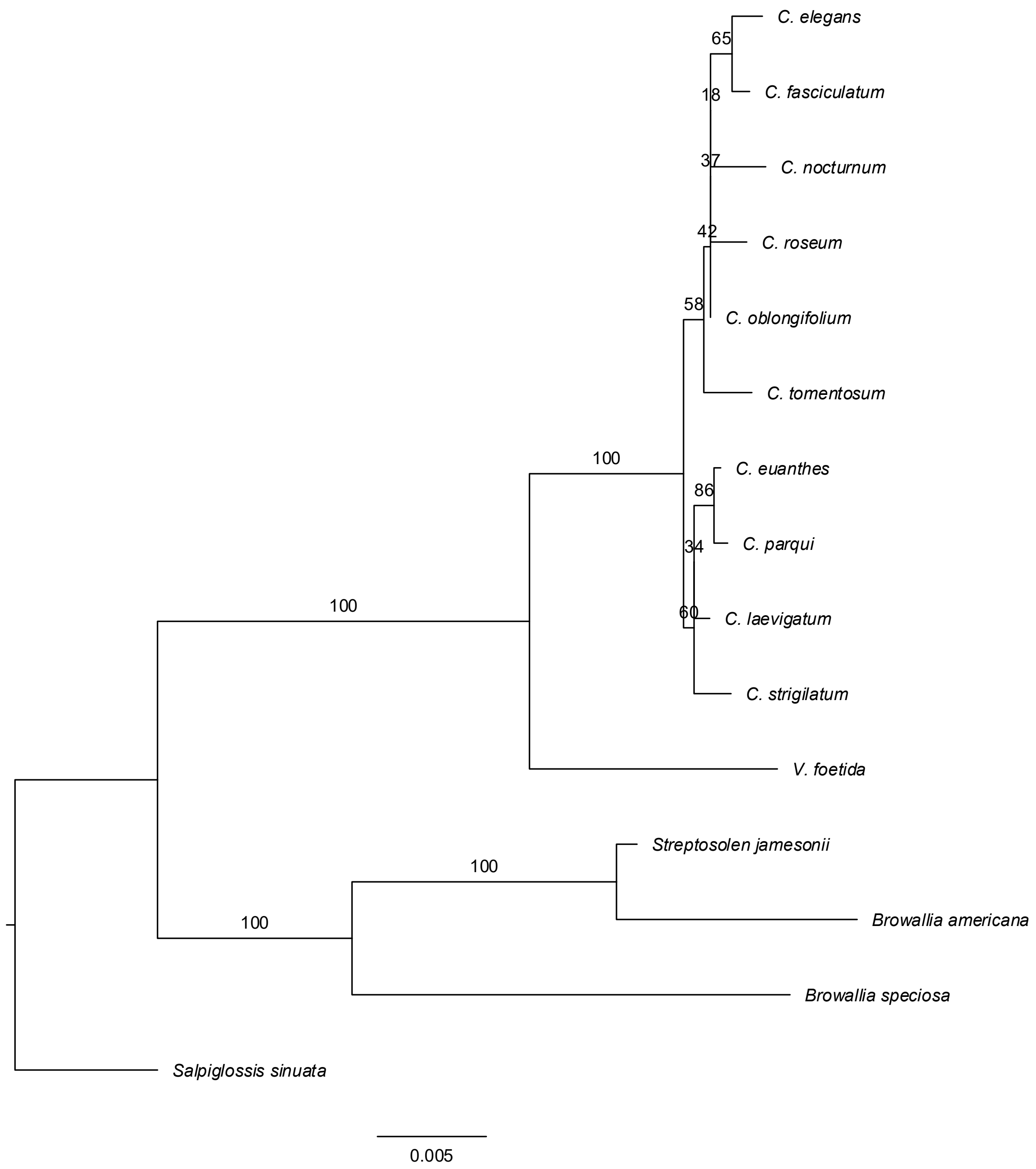


Supplementary Figure S3: ML analysis with *mat*K, *ndh*F and *trn*L-F.

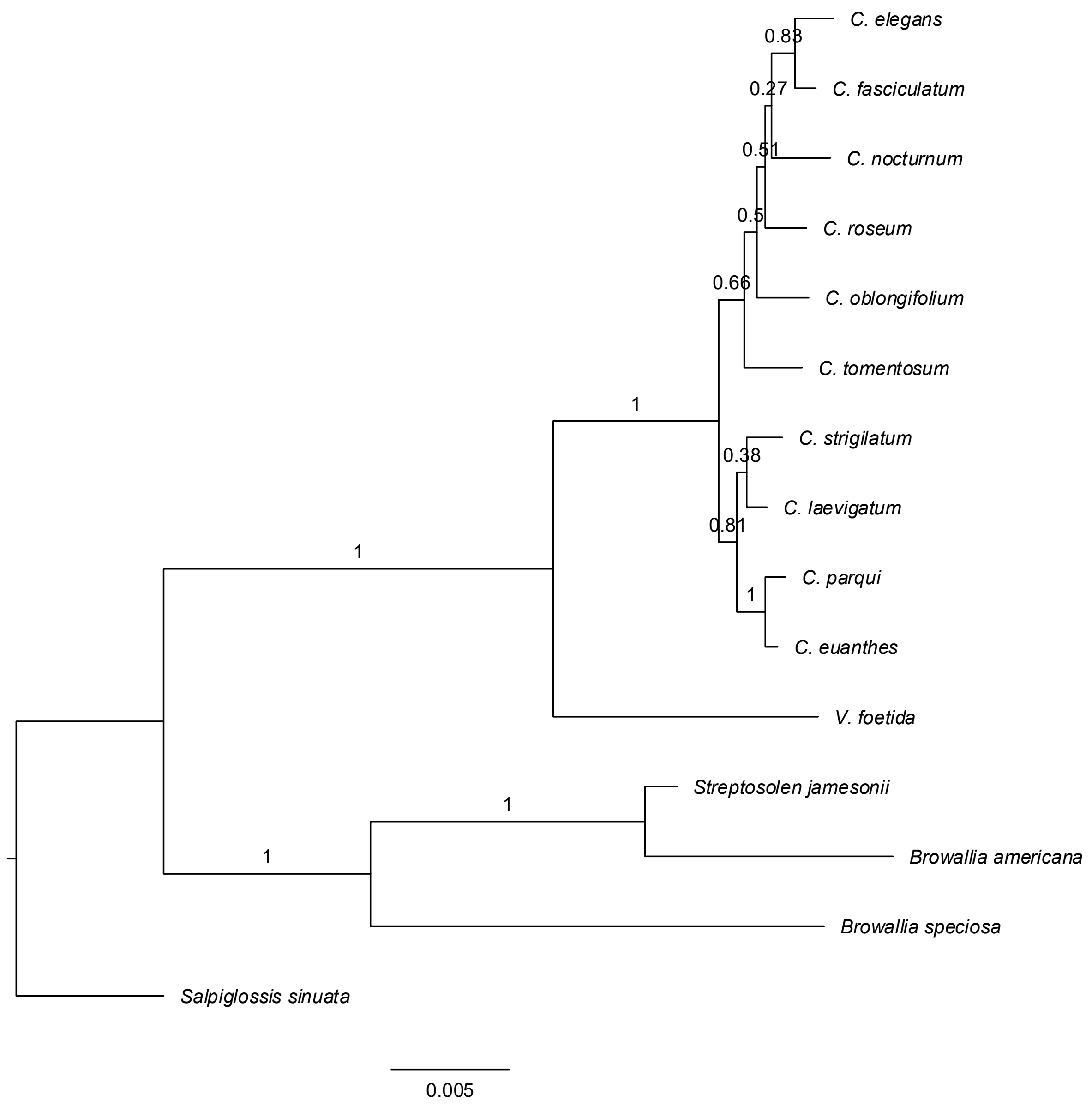


Supplementary Figure S4: BI analysis with *mat*K, *ndh*F and *trn*L-F

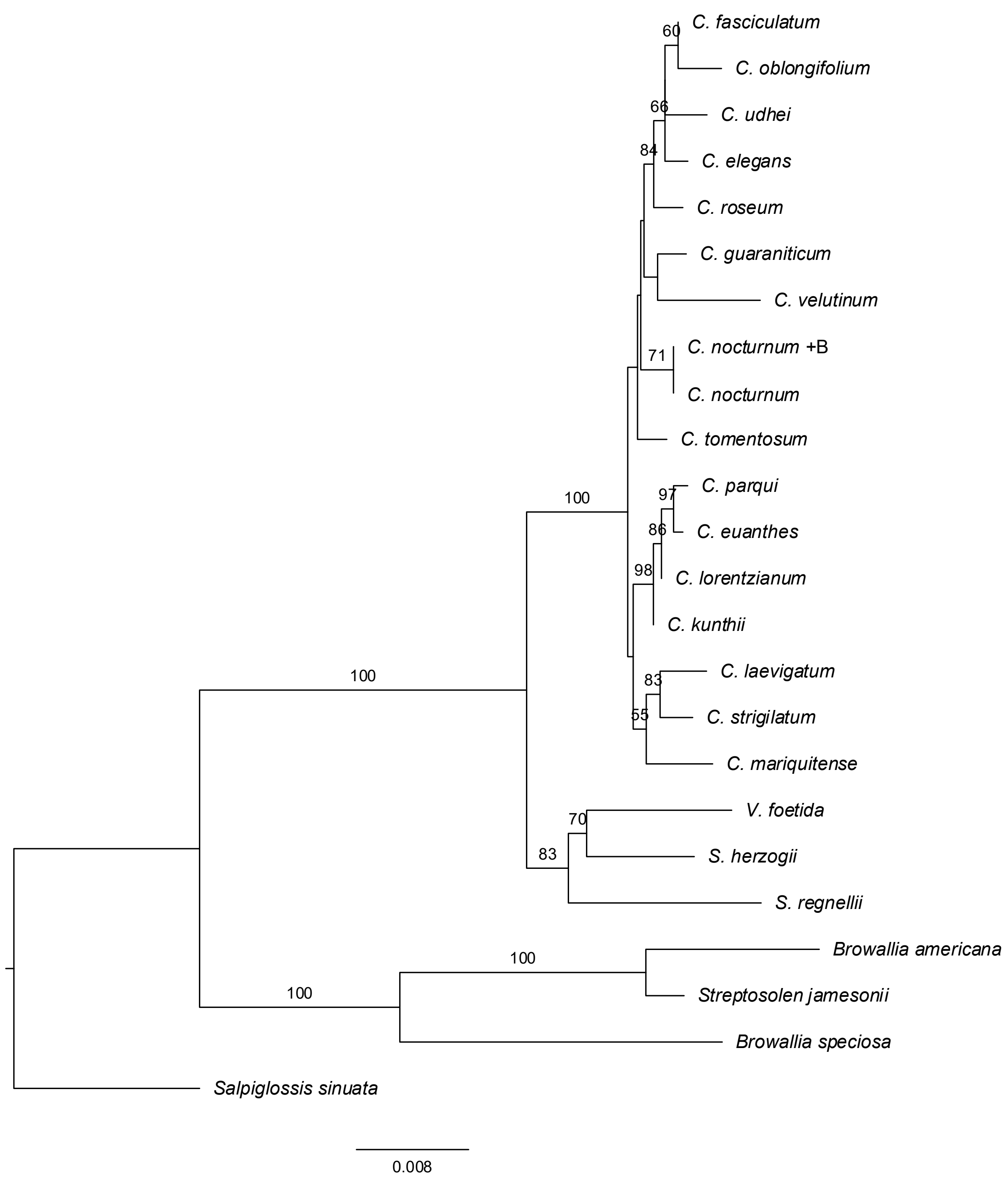


Supplementary Figure S5: ML analysis with ITS, *mat*K, *ndh*F and *trn*L-F.

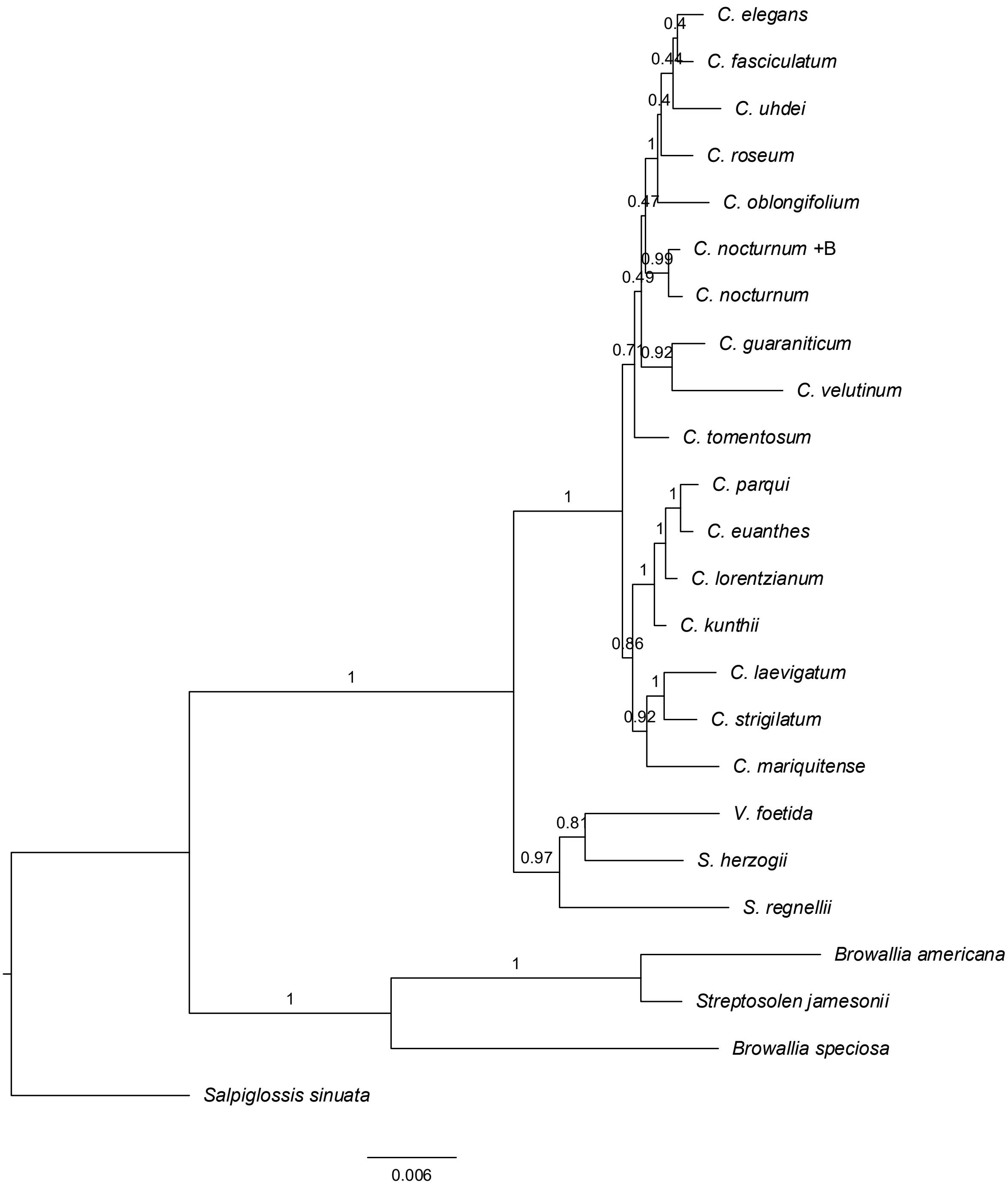


Supplementary Figure S6: BI analysis with ITS, *mat*K, *ndh*F and *trn*L-F.
