## Supplementary Table for "Phylogeny and chromosomal differentiation in Cestreae (Solanaceae)"

**Supplementary Table 1.** GenBank sequences.

| **Species** | **ITS** | **matK** | **ndhF** | **trnLF** |
| --- | --- | --- | --- | --- |
| *Cestrum elegans* | AJ492459 | AJ585891.1 | AJ585952.1 | KP756701.1 |
| *Cestrum euanthes* | PQ605139* | KP756848.1 | KP756890.1 | MG851859.1 |
| *Cestrum fasciculatum* |  | KP756849.1 | KP756892.1 | DQ508627.1 |
| *Cestrum guaraniticum* | PQ605140* |  |  |  |
| *Cestrum kunthii* | PQ605141* |  |  |  |
| *Cestrum laevigatum* | PQ605142* | MG875396.1 | KU376254.1 | MG851868.1 |
| *Cestrum lorentzianum* | PQ605143* |  |  |  |
| *Cestrum mariquitense* | PQ605144* |  |  |  |
| *Cestrum nocturnum +B* | PQ605146* |  |  |  |
| *Cestrum nocturnum* | PQ605145* | MF350140.1 | AY206741.1 | AY206723.1 |
| *Cestrum oblongifolium* | DQ508674 | DQ508571.1 |  | DQ508639.1 |
| *Cestrum parqui* | PQ605147* | MG875406.1 | KU376250.1 | MG851880.1 |
| *Cestrum roseum* | DQ508678 | MG875409.1 | KP756864.1 | DQ508643.1 |
| *Cestrum strigilatum* | PQ605148* | KP756810.1 | EU126008.1 | EU580976.1 |
| *Cestrum tomentosum* | DQ508682 | OQ289892.1 | EU126009.1 | EU580977.1 |
| *Cestrum uhdei* | PQ605149* |  |  |  |
| *Cestrum velutinum* | PQ605150* |  |  |  |
| *Sessea herzogii* | PQ605151* |  |  |  |
| *Sessea regnellii* | PQ605152* |  |  |  |
| *Vestia foetida* | DQ508652 | EF438822.1 | AY206751.1 | AY206769.1 |
| *Salpiglossis sinuata* | KP100299.1 | EF439055.1 | U08928.1 | KM200028.1 |
| *Browallia americana* | MK412104.1 | EF439050.1 | KU678197.1 |  |
| *Browallia speciosa* | KP100284.1 | KP756836.1 | AY206739.1 | AY206753.1 |
| *Streptosolen jamesonii* | KP100275.1 |  | EU580948.1 | EU581064.1 |

(*) The newly obtained sequences

**Supplementary Table 2.** AIC and deltaAIC values obtained from model comparison (jmodelTest) to be used in ML analysis.

| **Model** | **AIC** | **delta AIC** |
| --- | --- | --- |
| TVM+G | 24.968.354 | 0.0 |
| GTR+G | 24.968.526 | 172 |

**Supplementary Table 3.** AIC values obtained from the comparison of models used for the optimization of the ChromEvol.

| **Model** | **AIC** |
| --- | --- |
| CONST_RATE | 28.87 |
| CONST_RATE_DEMI | 30.25 |
| CONST_RATE_DEMI_EST | 35.73 |
| CONST_RATE_NO_DUPL | 64.66 |
| LINEAR_RATE | 32.92 |
| LINEAR_RATE_DEMI | 34.32 |
| LINEAR_RATE_DEMI_EST | 39.97 |
| LINEAR_RATE_NO_DUPL | 58.05 |
| BASE_NUMBER | 32.87 |
| BASE_NUMBER_NO_DUPL | 30.85 |

**Supplementary Table 4.**  AIC values obtained from the comparison of models used to Reconstruct the Ancestral States of the karyotypic formula and the rDNA distribution.

| **Model** | **Karyotype Formula** | **rDNA Distribution** | **rDNA Types** |
| --- | --- | --- | --- |
| ER | 16.429 | 59.288 | 46.698 |
| Direccional | 17.201 | 65.907 | 67.060 |
| Ordered | 19.857 | 65.907 | 68.478 |
| SYM | 19.125 | 55.397 | 54.042 |
| ARD | 23.764 | 65.907 | 81.257 |
